# The accuracy of *Apis mellifera* reference genomes: from version Amel 3.0 to Amel 4.5

**DOI:** 10.1101/2023.10.22.563459

**Authors:** Benjamin H. Conlon, Eike Oertelt, Jarkko Routtu

**Author notes:** **Corresponding author:** Benjamin H. Conlon, Current address: Molecular Ecology, Institute of Biology/Zoology, Martin-Luther-University Halle-Wittenberg, Hoher Weg 4, 06099 Halle an der Saale, Germany., Correspondence and requests for materials should be addressed to B.H.C. **Footnotes** Author contributions: data were analysed by B.H.C., the R phasing package was written by E.O., J.R. supervised the project. All authors contributed to the manuscript. The authors declare no competing financial interests.

## Abstract

The availabilty of reference genomes is accelerating rapidly, making their use in a wide variety of biological research programmes more feasible than ever. However, current Next-Generation Sequencing platforms are limited in the length of reads they are able to produce; requiring the correct order to be determined algorithmically. While there is a potential for errors in assembly algorithims, genetic pedigree data can be used to identify recombination events and, as recombination events are rare locally, test the order of sequences within a genome assembly. We use high-resolution population genomic data to test and compare the assembly quality of the three most recent reference genome assemblies for the western honey bee (*Apis mellifera*). As a model organism, there are several reference genomes available for *A. mellifera* with estimated recombination rates ranging from 19 cM/Mb to 37 cM/Mb. We identify a large degree of variation between assemblies and find that at least 20% of the most recent *A. mellifera* reference genome is mis-assembled. Providing an explanation for the degree of variation in estimated recombination rates and potentially influencing results downstream.

## Introduction

The prevalence of reference genomes^1,2,3,4^ makes sequencing a sub-sample of a genome more feasible for a wider range of experiments. Large volumes of genomic data can now be generated, aligned to a reference genome, and filtered for features of interest^5,6,7^. The process is both less labour intensive than screening using Polymerase Chain Reactions^7,8,9,10,11^ and less computationally intensive than *de novo* genome assembly^2,5^. However, downstream processing of the results relies on the assumption that the reference genome is correct. The short read lengths produced by current Next Generation Sequencing (NGS) platforms and the prevelance of repeat regions means that reference genomes are made up of distinct scaffolds and contigs rather than contiguous chromosomal sequences. These scaffolds are assembled algorithmically into chromosomes^12^, presenting a potential for errors^2^.

The error rate in the algorithmic assembly of a reference genome can be reduced by pairing it with population genetic data or experimental crosses^11,13,14,15,16,17,18^. Using SNP or microsatellite markers to identify recombination events, a genetic map can be created for the genome. The decay of linkage disequilibrium with increased physical distance can then be used to identify incorrectly-located regions within the reference genome and estimate their true location^17^.

As one of the first species identified for Whole Genome Sequencing (WGS)^15^, the western honey bee (*Apis mellifera*) has four available reference genome assemblies with both *de novo* and reference-genome-based genetic maps (Table 1)^11,12,14^. As haploid males can be analysed this makes it an excellent candidate model system for testing the reproducibility of genomic assemblies. While previous genetic maps, constructed *de novo* based on recombination frequency between marker positions in the scaffolds of the reference genome, have consistently reported a recombination rate of 19-22 cM/Mb (Table 1.), the most recent version of the genome (Amel_4.5)^12^ has provided estimates with much higher variability (Table 1.). Unusually, given the amount of previously-published population genetic data for *A. mellifera*, and its utility in previous genome assemblies^11,14,15^, no population genetic data was used in the construction of the Amel_4.5 reference genome^12^. This raises the possibility that large increases in recombination rate seen in genetic maps constructed using Amel_4.5 are due to assembly errors rather than genuine recombination events.

**Table 1.** Comparison of recombination rate estimates, for previous *de novo* and reference-genome-based genetic maps, to our estimate.

| Paper | Data-type | Genome assembly version | Recombination rate (cM/Mb) |
| --- | --- | --- | --- |
| Beye *et al.*, 2006 | Microsatellite | 2 | 19.0 |
| Solignac *et al.*, 2007 | Microsatellite | *de novo* | 22.0 |
| Wahlberg *et al.*, 2015 – did not estimate between scaffolds | Genomic | 4.5 | 26.0 |
| Liu *et al.*, 2015 – Upper estimate | Genomic | 4.5 | 37.0 |
| Liu *et al.*, 2015 – ignoring multiple recombination events | Genomic | 4.5 | 24.5 |
| Conlon *et al.* – 3.0 | Genomic | 3.0 | 42.2 |
| Conlon *et al.* – *de novo* | Genomic | 3.0 | 22.8 |
| Conlon *et al.* – 4.0 | Genomic | 4.0 | 38.5 |
| Conlon *et al.* – *de novo* | Genomic | 4.0 | 22.6 |
| Conlon *et al.* – 4.5 | Genomic | 4.5 | 98.1 |
| Conlon *et al.* – *de novo* | Genomic | 4.5 | 44.0 |

Given its importance as a model organism, we seek to test the variation in quality among the three most used genome assemblies for *A. mellifera*. The haplodiploid sex determination, found in *A. mellifera* and throughout the Hymenoptera, greatly simplifies the task of identifying recombination events as drones possess only one allele per locus: removing any ambiguities associated with heterozygosity. This, combined with a wide range of available genetic data, means *A. mellifera* is unusually well suited for a study of this kind.

## Results

### Genotyping results

After filtering in r/qtl, our datasets contained: 49 individuals with 1456 unique markers and 92% coverage for the 3.0 genome assembly; 48 individuals with 1556 unique markers and 92% coverage for the 4.0 genome assembly and 78 individuals with 2879 unique markers and 77% coverage for the 4.5 genome assembly.

### Map construction

Based on marker order in the reference genome, we calculated the recombination rates of 42.2 cM/Mb, 38.5 cM/Mb and 98.1 cM/Mb for the 3.0, 4.0 and 4.5 reference genome assemblies respectively. These are much higher than previous estimates using *de novo* assemblies (Table 1.). We identified high genetic linkage between physically distant markers, a sign of assembly errors, in all genome assemblies (S1), however, the 4.5 genome assembly is much worse (S1; S2); as evidenced by its recombination rate being more than double that of 3.0 or 4.0.

Having performed *de novo* genetic map constructions, the quality of each assembly improves (S1; S2). By comparing marker position in the genome assembly to the *de novo* assembly, we can see that much of the improvement has come from re-orientation of large regions within the linkage groups rather than a total rearrangement of markers (Figure 1 A, B, C). The estimated recombination rates for the new assemblies are 22.8cM/Mb and 22.6cM/Mb for the 3.0 and 4.0 genome assemblies respectively and 44cM/Mb for the 4.5 genome assembly. The estimated recombination rate for the *de novo* assemblies using markers form the 4.0 and 3.0 assemblies are very close to earlier estimates using *de novo* constructions (Table 1.). While the estimate for markers from the 4.5 genome assembly is higher than has previously been estimated (Table 1.), this may be a result of the poor assembly quality affecting the phasing script used.

**Figure 1.**
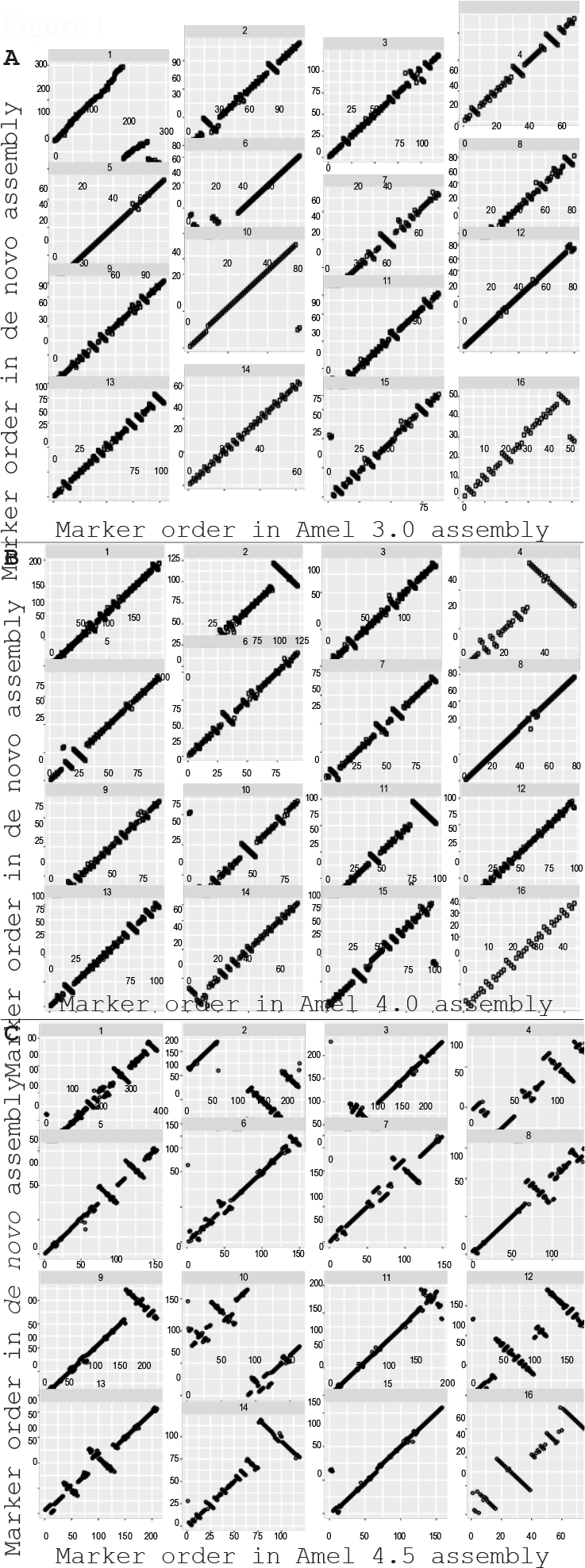
Comparison of marker order in the genome assembly vs the *de novo* map construction for (A) Amel_3.0 (n = 1456), (B) Amel_4.0 (n = 1556) and (C) Amel_4.5 (n = 2879).

### Relation to scaffold order

We identified 72 clearly misplaced or inverted regions within the current *Apis mellifera* 4.5 reference genome assembly (Figure 1 C; S3). Of these, 71 contained all the markers on a scaffold suggesting an error during genome construction. For the one region which did not contain all the markers on a scaffold, there was a large amount of variation in placement between the scaffold’s remaining markers suggesting a degree of error in the construction and that the entire scaffold is also incorrectly orientated.

## Discussion

We compared the genome construction quality among the three most recent versions of the *Apis mellifera* genome assembly using high-density SNP-based linkage maps. After performing *de novo* map constructions, we found the linkage between adjacent markers increased greatly while the number of recombination events decreased. As linkage is expected to decrease with increasing physical distance^17^, this supports our re-ordering of markers in the *de novo* map construction. For *Apis mellifera* 3.0 and 4.0, the two most well-constructed genome assemblies in our analysis, this results in recombination rates of 22.8cM/Mb and 22.6cM/Mb, highly similar to the 22.0cM/Mb reported in a previous *de novo* assembly^11^. While the estimated recombination rate for the *Apis mellifera* 4.5 genome assembly is higher (44cM/Mb), much of this difference could be explained by the misplacement or misorientation of at least 20% of scaffolds in the Amel_4.5 reference genome and the effect this would have on the phasing method used.

Despite being the current representative genome for *A. mellifera*, the Amel_4.5^12^ genome assembly was the worst constructed in our analysis; a result which could cause problems for the 86 studies which cite it in Web of Science (Clarivate Analytics). The lack of new population genetic data for the Amel_4.5 genome assembly^12^ means assembly quality was not experimentally tested and likely contributes to our identification of such a high number of mis-placed and mis-aligned scaffolds.

While *A. mellifera* does appear to have one of the highest recombination rates of any eukaryote^11,14,15,19^, the very high estimates associated with the Amel_4.5 genome assembly^19^ are likely biased by poor genome construction rather than representing genuine recombination events. Indeed, when studies do report recombination frequencies under 30 cM/Mb for the Amel_4.5 genome assembly (Table 1.), they have either ignored multiple recombination events between markers^19^ or did not estimate recombination events between scaffolds^20^: supporting our conclusion that many scaffolds are misplaced.

The generation of reference genomes is accelerating rapidly. On the 1^st^ of May 2017, 4991 (for 4314 unique species) eukaryotic genome assemblies were stored in the NCBI genome assembly database^21^. 1334 (27%) of these assemblies were added in 2016, at an average rate of over 111 per month; compared to a total of 50 (1%) genome assemblies added in the first five years and 403 (8%) in the first decade of this millenium^21^. While the rapid rise and proliferation of genomic data will benefit research, our results show that even well-resolved assemblies cannot be relied on fully and that high-density linkage maps generated using NGS can be invaluable testing assembly quality. The variation we find between reference genome assemblies also highlights the difficulties that come with comparing results to those generated using different genome assemblies.

The availability of reference data does make genome-level studies cheaper and more feasible for a wide range of studies^1,2,7^. However, caution should be exercised when using them and the quality of the construction should, if possible, be confirmed experimentally^11,13,14,15,16,17,18,22^. This would not only provide more reliable results for a single study but, by maximising the accuracy of each iteration of a reference genome assembly, should make comparisons between iterations more feasible than currently appears possible for *A. mellifera*.

## Materials and Methods

### Sampling and DNA analyses

We sampled drone offspring from a hybrid queen^10^, extracting DNA with a phenol/chloroform protocol^23^. The resulting extracts were assessed using a Nanodrop 1000 spectrophotometer (peqlab). Specimens were split into three sequencing runs: one of 16 samples and two of 32 samples. Library preparation was conducted using the RESTseq method^24^ and sequenced using IonTorrent (Thermo Fisher).

### SNP identification

Sequencing quality was checked using FastQC^25^ and barcodes trimmed with cutadapt^26^. The resulting sequences were mapped to the *Apis mellifera* 4.5, 4.0 and 3.0 genome assemblies^12,15^ using the “MEM” algorithm of the Burrows-Wheeler Aligner (BWA)^27^. The mapped reads were aligned using the reference genome with SAMtools’ mpileup function^28^ before variant loci were identified using BCFtools’ call function^29^.

### Defining of the Phase

The phase for each locus in a linkage group was determined by using an algorithm written in R^30^ (https://github.com/EikeOertelt/PhaseHaploid). The algorithm is based on the fact that recombination events are rare locally when using high density markers such as SNPs produced by RESTseq.

By identifying a shared locus between datasets, we were able to use this as an anchor point, allowing multiple datasets to be phased and then combined. Starting at the anchor point, each marker matching the reference genome was assigned the phase “A” and each marker different to it was assigned phase “B”. To overcome the problem that the expression (matching or different to the reference genome) could switch without the phase switching as well, we used precedents for each of the phases. A precedent for the “matching reference genome” phase would be a match in the expression of the current marker and the previous marker in the same individual. In case of a mismatch this would be a precedent for the “alternative to reference genome” phase. This was done for each linkage group. After working through all the linkage groups, the datasets were simply merged by the position in the genome.

### Map construction

Data were filtered with r/qtl^30,31^ to remove duplicate markers and individuals with under 1000 markers. Using the ASMap package in R^30,32^, recombination events and genetic distances were calculated for marker orders based on the reference genome assemblies before *de novo* map constructions were performed. Optimal marker order within a linkage group was determined by minimising the number of crossovers and genetic distances were calculated using the Kosambi map function. Due to changes in the marker order, phasing was checked and manually adjusted before re-running the *de novo* map construction. We then compared the physical location of incorrectly-placed regions to the reference genome structure.

## Acknowledgements

We would like to thank Petra Leibe and Denise Kleber for their help with the lab work, funding was provided by Deutsche Forschungsgemeinschaft grant RO 5121/1-1 to JR.

## Data deposition

Sequence data has been deposited in the Sequence Read Archive (SRA) of the National Centre for Biotechnology Information (NCBI) under the accession number: PRJNA388299

## Supplementary material

**S1.**
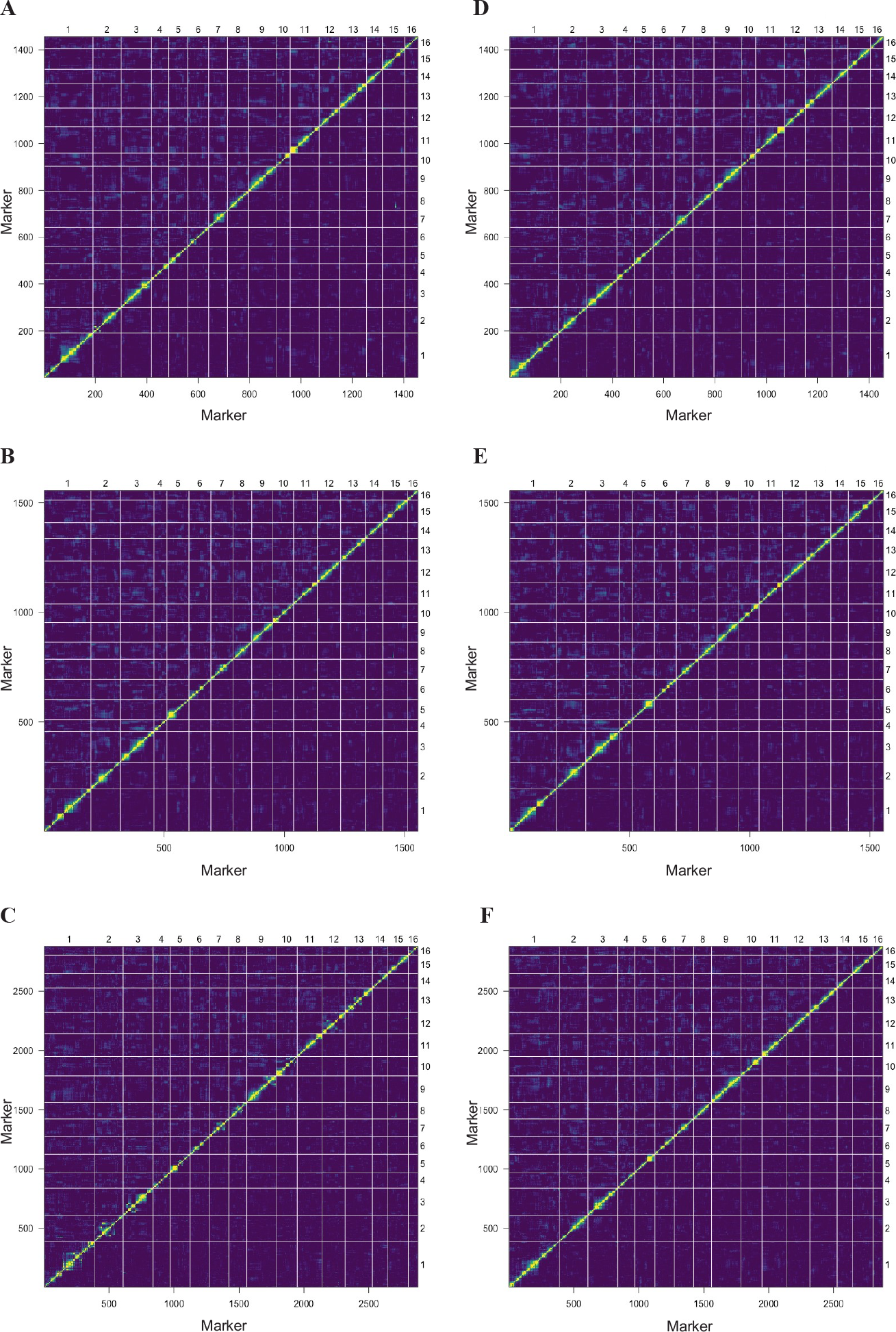
Heatmap showing plotting the pair-wise recombination fractions against the LOD linkage score for the (A) Amel_3.0 (n = 1556), (B) Amel_4.0 (n = 1556) and (C) Amel_4.5 (n = 2879) genome assemblies and the (D) *de novo* 3.0 (n = 1456), (E) *de novo* 4.0 (n = 1556) and (F) *de novo* 4.5 (n = 2879) maps. Yellow represents high genetic linkage and a low recombination rate between two markers while blue represents low genetic linkage and a high recombination rate between two markers.

**S2.**
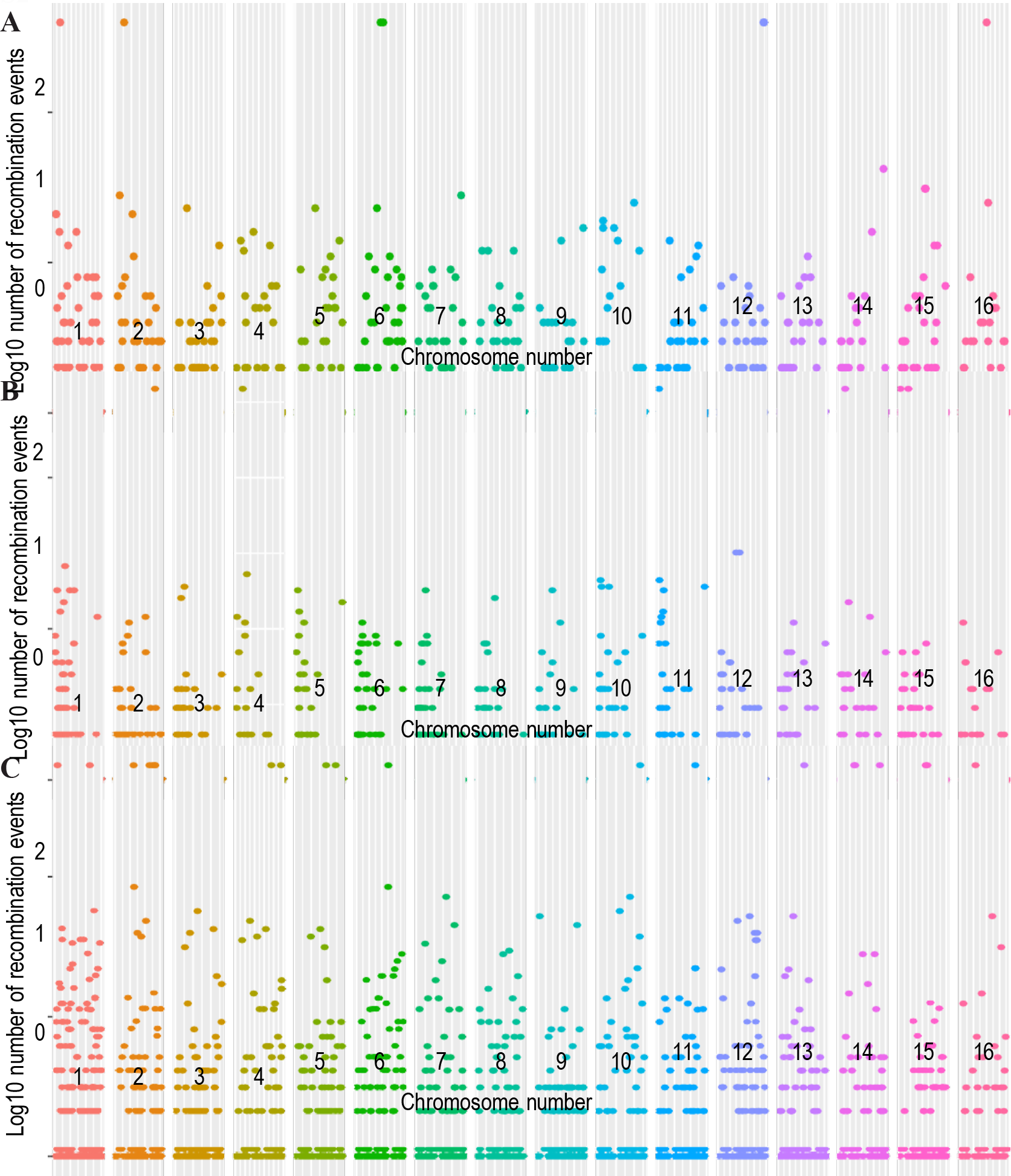
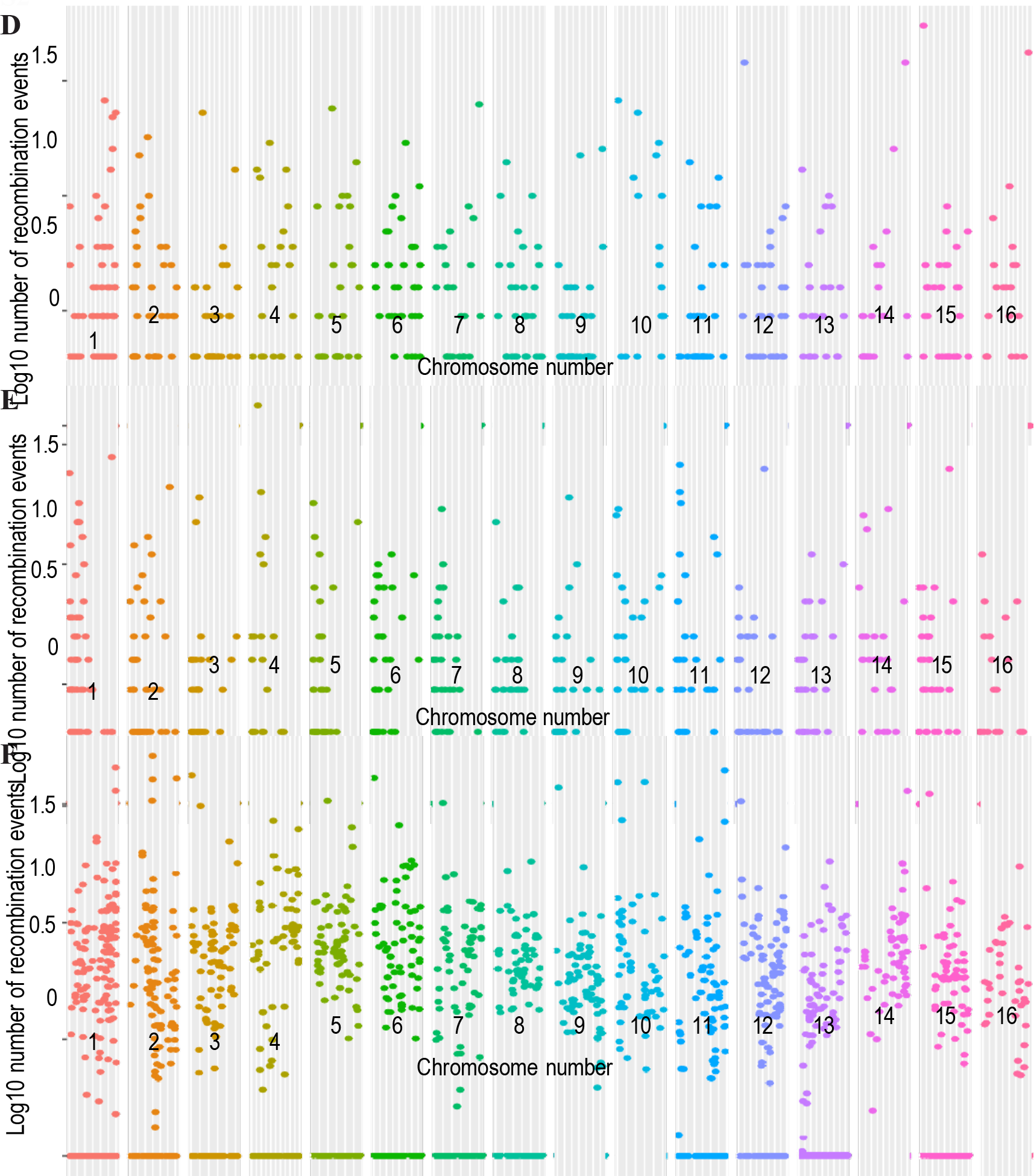
Log10 of recombination events between adjacent markers along the (A) Amel_3.0 (n = 1456), (B) Amel_4.0 (n = 1556) and (C) Amel_4.5 (n = 2879) genome assemblies and the (D) *de novo* 3.0 (n = 1456), (E) *de novo* 4.0 (n = 1556) and (F) *de novo* 4.5 (n = 2879) maps.

**S3.** Location of identified mis-aligned scaffolds in the *Apis mellifera* 4.5 reference genome assembly.

| Assembly | Chromosome | Upper region (Mbp) | Lower region (Mbp) | Upper Marker | Lower Marker | Scaffold | Entire Scaffold (Y/N) | Notes |
| --- | --- | --- | --- | --- | --- | --- | --- | --- |
| 4.5 | 1 | 17.79 | 16.12 | A212 | A184 | 29 | Y |  |
| 4.5 | 1 | 18.14 | 18.12 | A216 | A213 | 30 | Y |  |
| 4.5 | 1 | 18.73 | 18.37 | A224 | A217 | 31 | Y |  |
| 4.5 | 1 | 24.67 | 22.62 | A362 | A293 | 37 | Y |  |
| 4.5 | 1 | 29.79 | 28.56 | A482 | A444 | 43 | Y |  |
| 4.5 | 2 | 15.49 | 12.54 | A783 | A711 | 20 | Y | Misplaced |
| 4.5 | 2 | 8.42 | 7.47 | A647 | A574 | 15 | Y |  |
| 4.5 | 2 | 10.21 | 8.72 | A702 | A657 | 17 | Y |  |
| 4.5 | 3 | 6.12 | 4.12 | A910 | A834 | 8 | Y |  |
| 4.5 | 3 | 11.4 | 10.73 | A1017 | A1034 | 15 | Y |  |
| 4.5 | 4 | 5.42 | 5.02 | A1140 | A1130 | 8 | Y | Misplaced |
| 4.5 | 4 | 4.96 | 4.76 | A129 | A1127 | 7 | Y | Misplaced |
| 4.5 | 4 | 4.19 | 3.79 | A1126 | A1123 | 6 | Y | Misplaced |
| 4.5 | 4 | 1.73 | 2.98 | A1118 | A1082 | 5 | Y | Misplaced |
| 4.5 | 4 | 0.78 | 0.78 | A1079 | A1079 | 3 | Y | Misplaced |
| 4.5 | 4 | 1.2 | 0.93 | A1081 | A1080 | 4 | Y | Inverted and misplaced |
| 4.5 | 4 | 0.47 | 0.13 | A1078 | A1075 | 1 | Y | Misplaced |
| 4.5 | 4 | 5.65 | 7.4 | A1155 | A1142 | 9 | Y |  |
| 4.5 | 4 | 8.19 | 7.94 | A1159 | A1156 | 10 | Y |  |
| 4.5 | 4 | 11.58 | 9.38 | A1207 | A1167 | 13 | Y |  |
| 4.5 | 4 | 12.2 | 11.9 | A2108 | A1223 | 16 | Y |  |
| 4.5 | 5 | 8.56 | 7.65 | A1359 | A1327 | 12 | Y |  |
| 4.5 | 5 | 11.95 | 9.36 | A1412 | A1374 | 14 | Y |  |
| 4.5 | 6 |  |  |  |  |  |  |  |
| 4.5 | 7 | 6.26 | 5.42 | A1698 | A1692 | 17 | Y |  |
| 4.5 | 7 | 10.45 | 8.36 | A1791 | A1737 | 21 | Y |  |
| 4.5 | 8 | 8.58 | 7.17 | A1957 | A1917 | 9 | Y |  |
| 4.5 | 8 | 12.38 | 11.62 | A2015 | A1995 | 17 | Y |  |
| 4.5 | 9 | 2.96 | 2.82 | A2027 | A2026 | 7 | Y | Inverted and misplaced |
| 4.5 | 9 | 1.73 | 1.73 | A2023 | A2024 | 5 | Y | Inverted and misplaced |
| 4.5 | 9 | 11.07 | 9.63 | A2321 | A2223 | 12 |  |  |
| 4.5 | 10 | 12.84 | 6.03 | A2521 | A2414 | 23 – 29 | Y | Misplaced |
| 4.5 | 10 | 4.8 | 0.04 | A2412 | A2322 | 1 – 18 | Y | Misplaced |

S3
| Assembly | Chromosome | Upper region (Mbp) | Lower region (Mbp) | Upper Marker | Lower Marker | Scaffold | Entire Scaffold (Y/N) | Notes |
| --- | --- | --- | --- | --- | --- | --- | --- | --- |
| 4.5 | 11 | 14.38 | 13.87 | A2775 | A2742 | 20 | Y |  |
| 4.5 | 12 | 7.59 | 4.84 | A2901 | A2822 | 13 | Y |  |
| 4.5 | 12 | 8.91 | 8.23 | A2935 | A20910 | 16 | Y |  |
| 4.5 | 12 | 11.74 | 9.14 | A3016 | A2936 | 17 | Y |  |
| 4.5 | 13 | 2.13 | 1.49 | A3086 | A3052 | 4 | Y |  |
| 4.5 | 13 | 5.48 | 3.26 | A3169 | A3113 | 7 | Y |  |
| 4.5 | 14 | 0.56 | 0.36 | A3284 | A3280 | 1 | Y |  |
| 4.5 | 14 | 3.08 | 2.68 | A3314 | A3306 | 8 | Y |  |
| 4.5 | 14 | 8.39 | 7.71 | A3373 | A3361 | 14 | Y |  |
| 4.5 | 14 | 10.06 | 8.82 | A3421 | A3375 | 15 | Y |  |
| 4.5 | 16 | 0.24 | 0.01 | A3640 | A3642 | 1 | Y |  |
| 4.5 | 16 | 0.99 | 0.46 | A3643 | A3664 | 2 | Y |  |
| 4.5 | 16 | 4.09 | 2.32 | A3700 | A3667 | 4 | Y |  |
| 4.5 | 16 | 4.89 | 4.91 | A3704 | A3702 | 5 | Y |  |
| 4.5 | 16 | 5.74 | 5.49 | A3724 | A3716 | 6 | N |  |
| 4.5 | 16 | 7.06 | 6.24 | A3748 | A3727 | 9 | Y |  |

